# The astrocyte *Fabp7* gene regulates diurnal seizure threshold and activity-dependent gene expression in mice

**DOI:** 10.1101/2025.03.01.640632

**Authors:** Micah Lefton, Carlos C. Flores, Yuji Owada, Christopher J. Davis, Thomas N. Ferraro, Yool Lee, Wheaton L. Schroeder, Jason R. Gerstner

## Abstract

Epileptic seizures are often influenced by time-of-day and changes in vigilance state, yet the molecular and cellular mechanisms underpinning these associations remain poorly understood. Astrocytes, a pivotal type of glial cell, play a critical role in modulating neuronal excitability and circadian rhythms, and they express Fatty Acid Binding Protein 7 (Fabp7), a molecule vital for sleep regulation, lipid signaling, and gene transcription. This study investigates the role of *Fabp7* in determining time-of-day dependent seizure susceptibility. We assessed electroshock seizure thresholds in male C57/BL6N wild-type (WT) and *Fabp7* knockout (KO) mice. Results demonstrated that, compared to WT mice, *Fabp7* KO mice displayed significantly elevated general and maximal electroshock seizure thresholds (GEST and MEST) during the dark phase, but not during the light phase. To explore the impact of *Fabp7* on activity-dependent gene expression during seizures, we conducted RNA sequencing (RNA-seq) on cortical and hippocampal tissues from WT and *Fabp7* KO mice following MEST and SHAM procedures during the dark period. While immediate early genes (IEGs) showed considerable differential expression between WT-MEST and WT-SHAM, this expression was absent in *Fabp7* KO-MEST compared to *Fabp7* KO-SHAM. Gene ontology analyses revealed significant overlaps between the WT-MEST:WT-SHAM and *Fabp7* KO-SHAM:WT-SHAM comparisons, indicating that the basal mRNA expression profiles in *Fabp7* KO brains resemble those of WT brains in a post-ictal state. Collectively, these findings suggest that Fabp7 is a key regulator of time-of-day dependent neural excitability and that astrocyte-mediated signaling pathways involving Fabp7 interact with neuronal activity to influence gene expression in response to seizures.

**Significance Statement:** Changes in sleep/wake state and/or circadian time-of-day are thought to influence neural excitability, which may confer seizure susceptibility. Here we describe a role for astrocytic Fabp7 in regulating nocturnal seizure threshold and gene expression associated with differences in seizure susceptibility, introducing an astrocyte factor that may represent a novel antiepileptic target for drug development.

## Introduction

Epilepsy is a neurological disorder affecting around 65 million people worldwide, and studies over the past two decades have revealed that glial cells may play a crucial role in seizure etiology with strong therapeutic potential for treatment (1, 2). About 90% of drug-resistant epilepsy patients exhibit a circadian regulation of seizures, independent of the epilepsy type or brain region, which may also be influenced by sleep/wake behavior (3). Astrocytes, a type of glial cell, are known to affect neural excitability, changes in sleep/wake state and circadian rhythms (4); however, how these systems functionally interact to influence seizure susceptibility remain largely unknown. Recently, pathways regulating lipid-accumulated reactive astrocytes were shown to promote disease progression in epilepsy (5). Our previous studies have revealed that the astrocyte-enriched lipid binding protein, Fatty acid binding protein 7 (Fabp7), is regulated by core circadian clock components (6, 7), has a synchronized oscillation in gene expression throughout the brain (8-10), and regulates sleep across phylogenetically disparate species, from flies, to mice, to humans (11-15). Given *Fabp7* gene expression was elevated in dendritic layers of hippocampus by kainate-induced seizures (16) and is regulated by the circadian clock factor BMAL1 (6), which is also known to influence seizure threshold (17), Fabp7 may represent a molecular node for integrating changes in sleep/wake state, circadian rhythms, and lipid metabolism in astrocytes with seizure propensity (18, 19). Here we characterize time-of-day differences in electroshock seizure threshold in WT and *Fabp7* KO mice, changes in cortical/hippocampal gene expression, and subsequent gene ontology (GO) and pathway analyses in cross comparisons between WT-SHAM, WT-MEST, *Fabp7* KO-SHAM, and *Fabp7* KO-MEST mice.

## Results

We first determined time-of-day dynamics in seizure threshold in WT and *Fabp7* KO mice. Our experimental design included measuring seizure threshold at two timepoints in the light phase (ZT4 and ZT8) and the dark phase (ZT18 and ZT20) for both genotypes (Fig. 1A). We observed a significant difference in seizure threshold in both generalized (GEST) and maximal (MEST) seizures (p = 0.0008 and p < 0.0001, respectively, One-Way ANOVA), with a time-of-day dependent change in GEST and MEST in *Fabp7* KO compared to WT (Fig. 1B and C).

**Figure 1.**
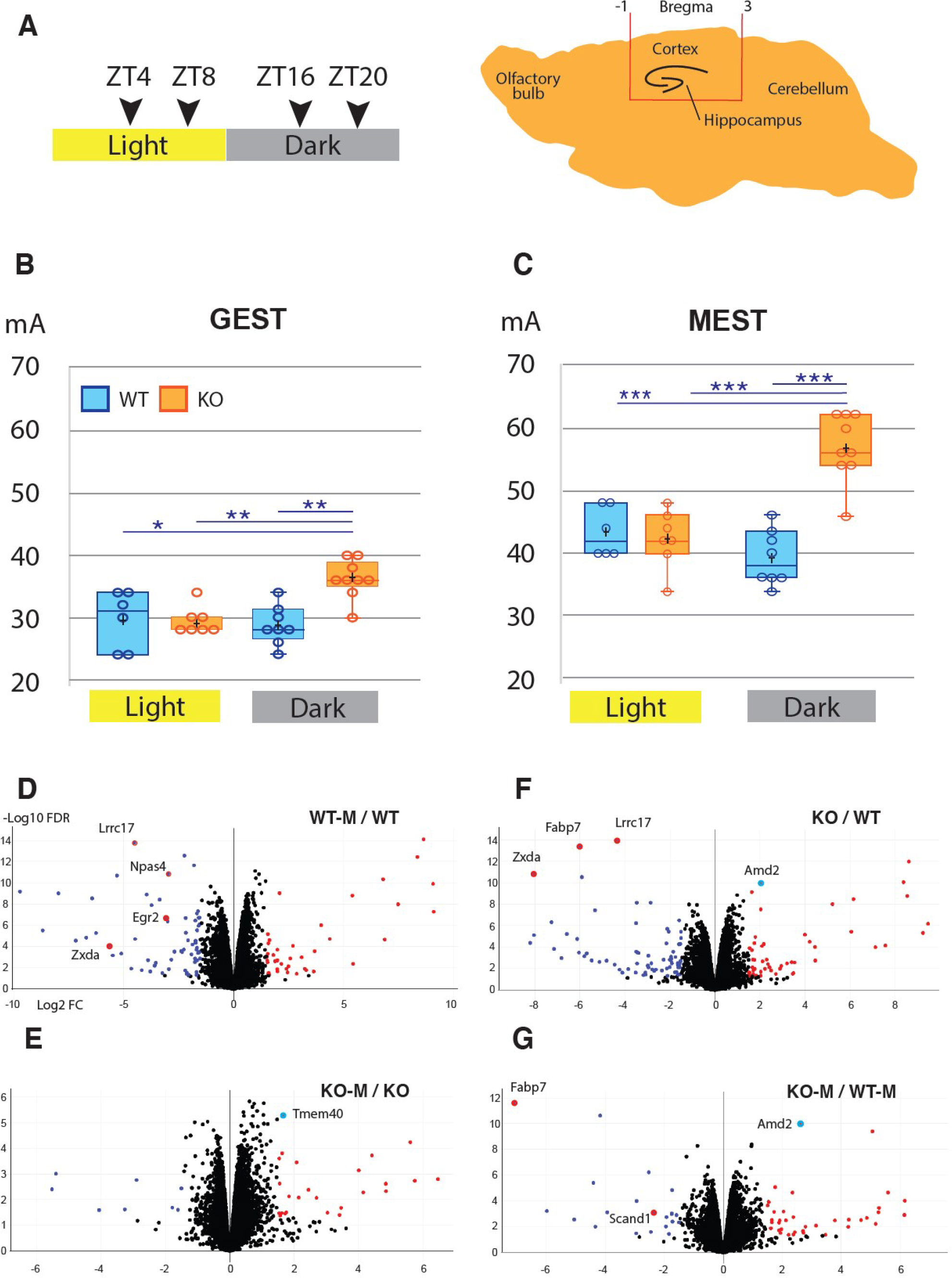
*Fabp7* KO mice have increased nocturnal seizure threshold associated with differential activity dependent gene expression compared to WT mice. (A) For each group of mice, electroshocks were delivered daily at specific times; ZT4 and ZT8 in the light period, and ZT16 and ZT20 in the dark period. At maximal seizure, brains were dissected, and the cortical/hippocampal region shown was collected from WT (N=5) and *Fabp7* KO (N=5) ZT20 mice for RNA-seq. (B) General and Maximal seizure thresholds for WT and KO mice during light and dark periods are plotted. Data for the two light timepoints and the two dark timepoints are combined: WT light (N=6), *Fabp7* KO-light (N=7), WT dark (N=8), *Fabp7* KO dark (N=9). One-way ANOVA, p = 0.0008 (GEST) and p < 0.0001 (MEST); post-hoc Bonferroni, *p<0.05, **p<0.01, **p<0.001. (D-G) Volcano plots of DEGs from RNA-seq results of cortical/hippocampal tissue for ZT20 MEST and SHAM mice. Log2 of fold change is plotted on the x-axis and - Log10 FDR is plotted on the y-axis.

As elevated seizure threshold levels in *Fabp7* KO versus WT mice were specific to the dark phase, we characterized alterations of bulk cortical/hippocampal tissue (Fig. 1A) transcriptomic signatures using RNA-seq during the nocturnal period for each genotype following recurring electroshock seizures and comparing them to control mice (SHAM) without seizures. Analysis of mRNA expression between WT-MEST and WT-SHAM identified many differentially expressed genes (DEGs; N=3857; FDR < 0.05), including several immediate-early genes (IEGs), such as *Npas4* and *Egr2* (Fig. 1D, *SI Appendix*, Dataset S1). However, when we evaluated *Fabp7* KO-MEST compared to *Fabp7* KO-SHAM, we did not observe nearly as many DEGs (N=201; FDR < 0.05), and many DEGs/IEGs identified in WT-MEST versus WT-SHAM were not affected by seizures in the *Fabp7* KO-MEST compared to *Fabp7* KO-SHAM mice (Fig. 1E, *SI Appendix*, Dataset S1). We then compared *Fabp7* KO-SHAM to WT-SHAM, which uncovered 2577 DEGs (FDR <0.05), including the genetic background control *Fabp7* mRNA (Fig. 1F). Unexpectedly, levels of DEGs in WT-MEST versus WT-SHAM (Fig. 1D, *SI Appendix*, Dataset S1) were similarly affected in *Fabp7* KO-SHAM to WT-SHAM, including *Lrrc17* and *Zxda* (Fig. 1F, *SI Appendix*, Dataset S1). Lastly, we compared *Fabp7* KO-MEST to WT-MEST, which identified 1570 DEGs (FDR <0.05), of which only few overlapped in *Fabp7* KO-SHAM versus WT-SHAM (e.g., *Fabp7* and *Amd2*; Fig. 1G, *SI Appendix*, Dataset S1).

GO and Pathway analysis between groups revealed RNA splicing and RNA binding were among the top overrepresented from downregulated genes between the WT-MEST versus WT-SHAM as well as the *Fabp7* KO-SHAM versus WT-SHAM comparisons, suggesting common transcriptional programming among these groups (Fig. 2, *SI Appendix*, Dataset S2). Overrepresentation among these two comparisons in upregulated GO and Pathways included protein targeting to membrane, ribosomal regulation, translation, and mitochondrial respiration and electron transport (Fig. 2). This similarity included neurodegenerative disease pathways, such as Parkinson’s, Huntington’s, and Alzheimer’s diseases. Remarkably, almost all of these shared overrepresented GO and Pathways between WT-MEST:WT-SHAM and *Fabp7* KO-SHAM:WT-SHAM were inversely overrepresented and shared between *Fabp7* KO-MEST:WT-MEST and *Fabp7* KO-MEST:*Fabp7*-SHAM groups (Fig. 2).

**Figure 2.**
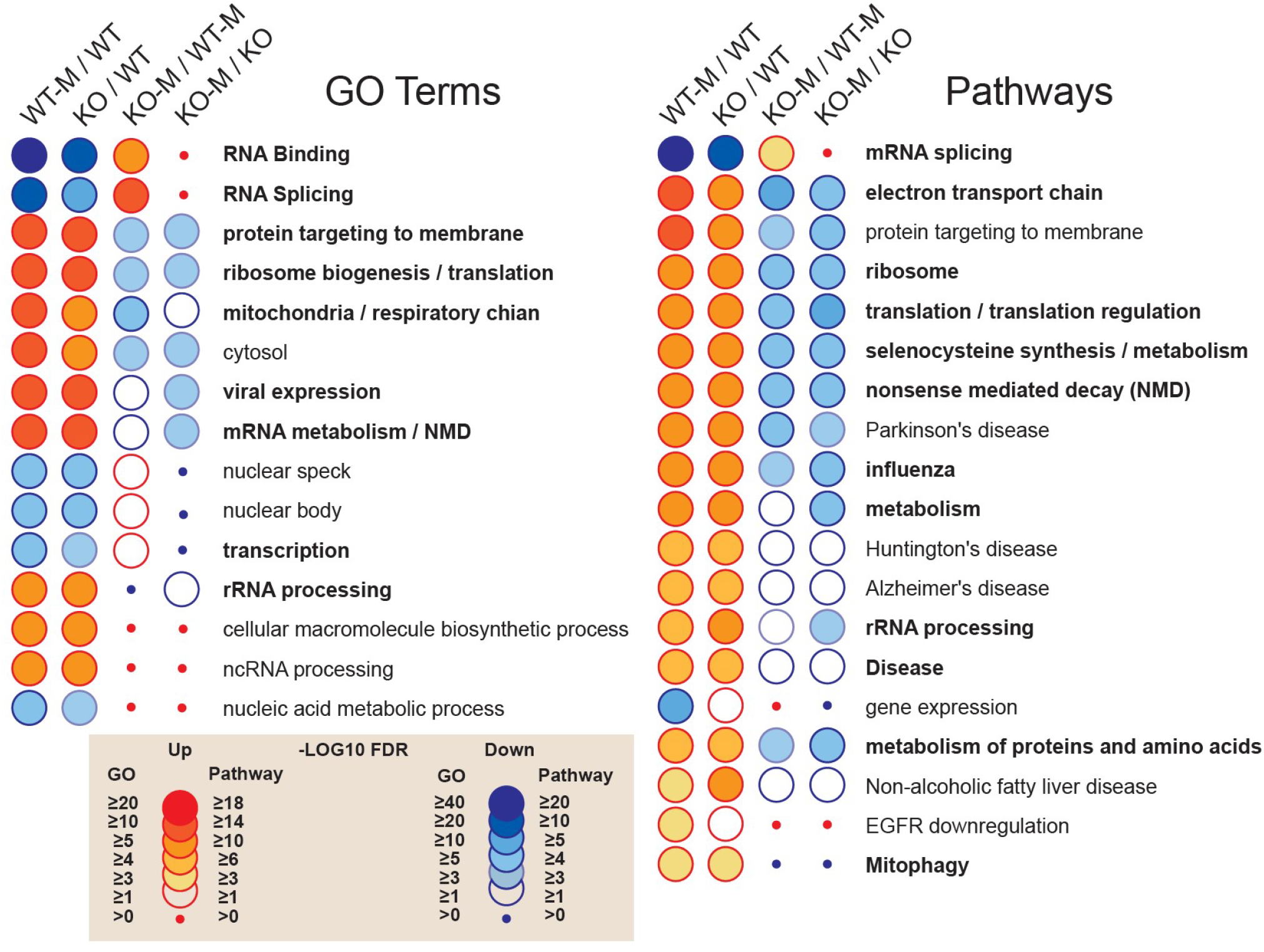
GO and Pathway analyses of DE genes in various comparisons between WT-MEST, WT-SHAM, KO-MEST and KO-SHAM. GO and Pathway overrepresentation reveal similar patterns between WT-MEST/WTSHAM and KO-SHAM/WT-SHAM. Many of these GO and Pathway patterns were reversed in the KO-MEST/WTMEST and KO-MEST/KO-SHAM comparisons. Threshold of the GO terms and Pathways is FDR <0.1.

## Discussion

We observed a significantly higher GEST and MEST in the nocturnal (wake) period of *Fabp7* KO compared to WT mice (Fig. 1B and C), which corresponded with disruption of seizure-associated DEGs (Fig. 1D-G, *SI Appendix*, Dataset S1) and their pathways (Fig. 2, *SI Appendix*, Dataset S2). Recently, local wake slow-wave (LoWS) activity showed progressive adaptive responses following network excitability prior to interictal epileptiform discharges (IEDs), which reduced the impact of subsequent IEDs (20). How these LoWS changes relate to variability in network seizure paths on circadian and slower timescales in patients with focal epilepsy remain to be determined (21). Comparing wake-dependent network activity relationships in electroencephalograph (EEG) signatures between *Fabp7* KO and WT mice before and after seizures merits future investigation, and may support an epilepsy homeostasis hypothesis, wherein LoWS reduces aberrant brain activity (20).

In summary, these results show that *Fabp7* is required for normal neural activity-dependent molecular and cellular processes and suggest that nocturnal astrocyte-regulated *Fabp7*-mediated signaling cascades are necessary for seizure responses. Given that about 30% of epilepsy patients eventually progress to a drug-resistant state (22), with glial scar formation and reactive glia at the epileptic focus involving astrocyte-derived lipid transport mechanisms (5, 23), Fabp7 may represent a novel therapeutic target to treat certain intractable forms of epilepsy.

## Materials and Methods

### Animals

All studies were approved by the Institutional Animal Care and Use Committees at the Washington State University (WSU; ASAF #6509) in accordance with the guidelines of the US National Institutes of Health. Experiments involved C57BL/6N wild type mice (The Jackson Laboratory, Bar Harbor, ME) and coisogenic Fabp7 knockout (KO) mice (from Y. Owada) and were bred in-house at the WSU Health Sciences Campus vivarium. Fabp7 KO mice were maintained as a homozygous strain. Litters were weaned between 21–22 days and pups were group housed by sex until the age of 13-16 weeks when they were entered into the study. Mice were maintained on a 12:12 hour light:dark cycle (lights on Zeitgeber time (ZT) 0, lights off ZT12) with access to food and water ad-libitum.

### Seizure tests and tissue dissection

Due to the variable effect of estrous cycle on seizure susceptibility (3), only male mice were studied. WT and Fabp7 KO mice (N=6-9 per group and condition) were tested for seizure threshold at 4 timepoints (ZT4, ZT8, ZT18, and ZT20), using a single electric shock delivered via ear clip electrodes once per day. SHAM mice received the same handling but did not receive the shock. We used a constant current electroshock unit (model No. 7801, Ugo Basile, Varese, Italy) in which the initial current level was set at 20 mA and increased by 2 mA with each successive daily trial until a maximal seizure, defined by bilateral tonic hind limb extension was observed. Other parameters of the stimulus were held constant (60 Hz, 0.4 ms pulse width, 0.2 s duration). Upon MEST, mice were euthanized, and brains were harvested immediately from WT-MEST (N=5), Fabp7 KO-MEST (N=5), WT-SHAM (N=5), and Fabp7 KO-SHAM mice (N=4) at ZT20 and flash frozen and kept at -80C until processing. Unilateral brain tissue encompassing the hippocampus and cortex (-1 to 3 mm AP and 0 to 2.5 mm ML) was blocked on dry ice, and homogenates used for subsequent RNA extraction, library construction, Illumina sequencing and data analysis.

### RNA isolation, cDNA synthesis, and high throughput sequencing

Total RNA was purified using the RNeasy Mini Kit (Qiagen). The integrity of total RNA was assessed using Fragment Analyzer (Advanced Analytical Technologies, Ankeny, IA) with the High Sensitivity RNA Analysis Kit. RNA samples with RQNs ranging from 8 to 10 were used for RNA library preparation with the TruSeq Stranded mRNA Library Prep Kit (Illumina, San Diego, CA). Briefly, mRNA was isolated from 1-2.5 µg of total RNA using poly-T oligo attached to magnetic beads and then subjected to fragmentation, followed by cDNA synthesis, dAtailing, adaptor ligation and PCR enrichment. The sizes of RNA libraries were assessed by Fragment Analyzer with the High Sensitivity NGS Fragment Analysis Kit. The concentrations of RNA libraries were measured using the StepOnePlus Real-Time PCR System (ThermoFisher Scientific, San Jose, CA) with the KAPA Library Quantification Kit (Kapabiosystems, Wilmington, MA). DNA was sequenced from both ends (paired-end) with a read length of 150 bp. The raw bcl files were converted to fastq files using the software program bcl2fastq, and adaptors were trimmed from the fastq files during the conversion. Sequence data (FASTQ files) were processed by trimming low-quality reads using Trimmomatic (version 0.39). and removing rRNA sequences using SortMeRNA (version 4.3.5). The remaining reads were aligned to the Mus musculus reference genome (mm10, UCSC) using HISAT2 (version 2.2.1). Gene expression quantification was analyzed using featureCounts (part of the Subread package, version 2.0.3).

### Data analysis

Generalized and maximal seizures were expressed as arithmetic mean values for each experimental group. One-Way ANOVAs were used to examine the effect of time of day on seizure thresholds. Post hoc analyses to examine statistical relationships for seizure threshold values between individual groups were conducted using the Bonferroni test (Prism stats package, v5). For analysis of seizure data, comparisons were collapsed to lights-on and lights-off phases since there were no significant differences between groups within each phase. Differential expression, volcano plots, GO and Pathway analysis of our data were conducted using Biojupies (https://maayanlab.cloud/biojupies/).

## Acknowledgments

The authors would like to thank Vivian Wei for technical assistance and the WSU-Spokane Genomics Core for expertise in RNA-sequencing.

## Data Availability

RNA-seq data are deposited in NCBI GEO (Accession #GSE271985).

## References

1. D. K. Binder, C. Steinhäuser, Astrocytes and Epilepsy. Neurochem Res 46, 2687–2695 (2021).

2. D. C. Patel, B. P. Tewari, L. Chaunsali, H. Sontheimer, Neuron-glia interactions in the pathophysiology of epilepsy. Nat Rev Neurosci 20, 282–297 (2019).

3. P. J. Karoly et al., Cycles in epilepsy. Nat Rev Neurol 17, 267–284 (2021).

4. M. H. Hastings, M. Brancaccio, M. F. Gonzalez-Aponte, E. D. Herzog, Circadian Rhythms and Astrocytes: The Good, the Bad, and the Ugly. Annu Rev Neurosci (2023).

5. Z. P. Chen et al., Lipid-accumulated reactive astrocytes promote disease progression in epilepsy. Nat Neurosci 26, 542–554 (2023).

6. J. R. Gerstner, G. K. Paschos, Circadian expression of Fabp7 mRNA is disrupted in Bmal1 KO mice. Mol Brain 13, 26 (2020).

7. W. M. Vanderheyden, B. Fang, C. C. Flores, J. Jager, J. R. Gerstner, The transcriptional repressor Reverbα regulates circadian expression of the astrocyte Fabp7 mRNA. Current Research in Neurobiology 2, 100009 (2021).

8. J. R. Gerstner, W. M. Vander Heyden, T. M. Lavaute, C. F. Landry, Profiles of novel diurnally regulated genes in mouse hypothalamus: expression analysis of the cysteine and histidine-rich domain-containing, zinc-binding protein 1, the fatty acid-binding protein 7 and the GTPase, ras-like family member 11b. Neuroscience 139, 1435–1448 (2006).

9. J. R. Gerstner et al., Brain fatty acid binding protein (Fabp7) is diurnally regulated in astrocytes and hippocampal granule cell precursors in adult rodent brain. PLoS One 3, e1631 (2008).

10. J. R. Gerstner et al., Time of day regulates subcellular trafficking, tripartite synaptic localization, and polyadenylation of the astrocytic Fabp7 mRNA. J Neurosci 32, 1383–1394 (2012).

11. J. R. Gerstner et al., Normal sleep requires the astrocyte brain-type fatty acid binding protein FABP7. Sci Adv 3, e1602663 (2017).

12. J. R. Gerstner, W. M. Vanderheyden, P. J. Shaw, C. F. Landry, J. C. Yin, Cytoplasmic to nuclear localization of fatty-acid binding protein correlates with specific forms of long-term memory in Drosophila. Commun Integr Biol 4, 623–626 (2011).

13. J. R. Gerstner, W. M. Vanderheyden, P. J. Shaw, C. F. Landry, J. C. Yin, Fatty-acid binding proteins modulate sleep and enhance long-term memory consolidation in Drosophila. PLoS One 6, e15890 (2011).

14. J. R. Gerstner et al., Amyloid-beta induces sleep fragmentation that is rescued by fatty acid binding proteins in Drosophila. J Neurosci Res (2016).

15. W. M. Vanderheyden, M. Lefton, C. C. Flores, Y. Owada, J. R. Gerstner, Fabp7 Is Required for Normal Sleep Suppression and Anxiety-Associated Phenotype following Single-Prolonged Stress in Mice. http://dx.doi.org/10.3390/neuroglia3020005.

16. Y. Owada, T. Yoshimoto, H. Kondo, Increased expression of the mRNA for brain- and skin-type but not heart-type fatty acid binding proteins following kainic acid systemic administration in the hippocampal glia of adult rats. Brain Res Mol Brain Res 42, 156–160 (1996).

17. J. R. Gerstner et al., BMAL1 controls the diurnal rhythm and set point for electrical seizure threshold in mice. Front Syst Neurosci 8, 121 (2014).

18. C. J. Re, A. I. Batterman, J. R. Gerstner, R. J. Buono, T. N. Ferraro, The Molecular Genetic Interaction Between Circadian Rhythms and Susceptibility to Seizures and Epilepsy. Front Neurol 11, 520 (2020).

19. J. R. Gerstner, C. C. Flores, M. Lefton, B. Rogers, C. J. Davis, FABP7: a glial integrator of sleep, circadian rhythms, plasticity, and metabolic function. Front Syst Neurosci 17, 1212213 (2023).

20. L. Sheybani et al., Wake slow waves in focal human epilepsy impact network activity and cognition. Nat Commun 14, 7397 (2023).

21. G. M. Schroeder et al., Seizure pathways change on circadian and slower timescales in individual patients with focal epilepsy. Proc Natl Acad Sci U S A 117, 11048–11058 (2020).

22. A. Fattorusso et al., The Pharmacoresistant Epilepsy: An Overview on Existent and New Emerging Therapies. Front Neurol 12, 674483 (2021).

23. P. Hayatdavoudi, M. Hosseini, V. Hajali, A. Hosseini, A. Rajabian, The role of astrocytes in epileptic disorders. Physiol Rep 10, e15239 (2022).

